# The RdRp Thumb-1 Pocket is a Conserved Target for Broad-Spectrum Antiviral Development

**DOI:** 10.1101/2024.03.29.587401

**Authors:** Virgil Woods, Tyler Umansky, Sean M Russell, Philippe Gallay, Davey Smith, Daniel Haders

**Affiliations:** Model Medicines, San Diego, CA, USA; Department of Immunology & Microbiology, Scripps Research, La Jolla, CA, USA; School of Medicine, University of California, San Diego, La Jolla, CA, USA

**Keywords:** RNA-dependent RNA polymerase, Thumb-1 allosteric site, broad-spectrum antivirals, hepatitis C virus, SARS-CoV-2, NS5B, nsp12, non-nucleoside inhibitors

## Abstract

RNA viruses cause human diseases ranging from mild colds to deadly pandemics. Direct-acting, broad-spectrum, non-nucleoside antivirals have been characterized as impossible to develop because allosteric binding sites are poorly conserved. The HCV NS5B RNA-dependent RNA polymerase (RdRp) Thumb-1 allosteric site and its interaction with the HCV NS5B Λ1-loop governs an essential conformational change required for polymerase initiation. The only approved NS5B Thumb-1 inhibitor, beclabuvir, has been shown to be inactive against a broad panel of non-HCV viruses, including poliovirus, rhinovirus, coronavirus, coxsackievirus, influenzavirus, and HIV. A conserved, homologous allosteric site on RdRp that spans multiple viral families has not been reported. Here, we report GALILEO’s discovery that the Thumb-1 pocket, its associated Λ1-loop and their interaction are conserved across RNA viral families for the first time. The discovery is structurally validated through comparative analysis of Protein Data Bank (PDB) deposited viral polymerases utilizing a method that allows independent, public validation. We further demonstrate that beclabuvir’s dependence on its indole C6 carbonyl to interact with the HCV-specific residue R^503^ restricts its activity to HCV. The discovery is biochemically validated with MDL-001, which does not contain a C6 carbonyl substituent. MDL-001 directly blocks viral RNA synthesis in isolated HCV and SARS-CoV-2 replication complexes and selects for the canonical Thumb-1 resistance mutation P495S in HCV NS5B. The discovery is validated at the cellular level through demonstration of MDL-001’s broad-spectrum, in vitro inhibition of both HCV and SARS-CoV-2. Preclinical proof of concept and development of MDL-001 across HCV, HBV, HDV, influenza, SARS-CoV-2, and RSV have been previously reported. These findings establish RdRp Thumb-1 as a conserved allosteric pocket and a druggable target for broad-spectrum direct-acting antiviral development.

## 1. Introduction

RNA virus pandemics and epidemics occur regularly and cause widespread death and economic devastation. In the last 25 years, humanity has endured at least six major viral outbreaks: West Nile Virus (1999–2002), SARS-CoV-1 (2002–2004), Swine Flu (2009–2010), MERS-CoV (2012-present), Ebola (2014–2016), and COVID-19 (2019-present) (1). The COVID-19 pandemic has caused an estimated 7 million deaths, though excess mortality analyses suggest the true toll lies between 19 and 36 million deaths (2). In two years, COVID-19 killed more people than any single outbreak since the 1918 influenza pandemic (2–4). These losses occurred despite unprecedented global coordination. The next pandemic may rise due to an established pathogen or a novel emergence within an existing viral family. Either way, the world is no better prepared for future pandemics than it was for previous ones (1, 4).

Respiratory viruses pose the greatest pandemic threat due to their transmission efficiency and capacity for sustained human-to-human spread following zoonotic emergence (5). Coronaviruses have caused three major outbreaks since 2002 and transmit repeatedly from animal reservoirs (5). Future pandemic threats are likely to emerge from the same family.

No direct-acting non-nucleoside antiviral has ever received regulatory approval for treatment of infections spanning multiple RNA viral families (1). Viral genomes are small and their replication strategies are divergent, leaving limited opportunities for cross-family targeting. The consequences in respiratory medicine are severe. Specifically, no approved antiviral treats more than one of the annual ‘tripledemic’ of influenza, Respiratory Syncytial Virus (RSV), and SARS-CoV-2 (6). RSV has no approved small-molecule therapy in adults. At the bedside, the absence of direct-acting broad-spectrum options forces clinicians to await diagnostic confirmation before initiating treatment. This clinical workflow often delays therapy beyond the narrow window during which existing options such as neuraminidase inhibitors remain effective (7–8). Infectious disease physicians urgently need a potent small-molecule direct-acting antiviral capable of rapidly treating viral infections immediately upon exposure or presentation of influenza-like illness (ILI) symptoms.

The novel COVID-19 pandemic exposed the cost of this therapeutic gap. No direct-acting broad-spectrum antivirals were stockpiled and ready for deployment against the novel SARS-CoV-2 virus. The direct-acting small-molecule options that later emerged have significant limitations. Remdesivir was the first drug shown to be clinically effective against SARS-CoV-2 (9). However, its intravenous route of administration confines its use to hospitalized patients, precluding the early outpatient treatment needed to maximize antiviral efficacy (10–11). Nirmatrelvir/ritonavir (Paxlovid) later provided an oral option. However, Paxlovid activity is limited to coronaviruses, with no activity against influenza, paramyxoviruses, or other respiratory pathogens (12). The current available treatment options leave humanity therapeutically as unprepared for the next novel pandemic as we were for the emergence of SARS-CoV-2.

Allosteric inhibition is established antiviral practice within individual viruses and individual viral families. The HCV NS5B polymerase carries four principal non-nucleoside sites (13). Thumb-1 and Thumb-2 sit on the thumb domain. Palm-1 and Palm-2 sit on the palm domain next to the active site. Each site has yielded clinical chemistry. Beclabuvir and deleobuvir bind Thumb-1, lomibuvir and filibuvir bind Thumb-2, dasabuvir binds Palm-1, and nesbuvir binds Palm-2 (13). Each of these compounds was developed for HCV; none are reported to treat another viral family. The same pattern holds outside HCV. The non-nucleoside inhibitor binding pocket of HIV-1 reverse transcriptase supports efavirenz, rilpivirine and doravirine. That chemistry does not inhibit HIV-2, a second species of the same genus (14). Baloxavir marboxil binds the influenza PA endonuclease and blocks cap-snatching, the step that initiates viral transcription. Its activity is confined to influenza A and B (15). Allosteric antiviral chemistry has therefore been understood to be virus or viral family-specific by construction.

The term “broad-spectrum” has historically been utilized to describe antiviral activity across viruses within a single family. Here we define “broad-spectrum” as antiviral activity across viral families, demonstrated at the structural, biochemical and cellular level. The development of a direct-acting broad-spectrum non-nucleoside antiviral has been considered impossible (16–17). Conventional wisdom holds that allosteric sites on viral polymerases are poorly conserved, restricting non-nucleoside inhibitors to genotype- or species-specific activity (13). The HCV NS5B RdRp Thumb-1 allosteric site provides a test of this dogma. Known inhibitors block an essential Λ1-loop mediated conformational change required for polymerase initiation (18) (**Fig. 1**). Beclabuvir is the only approved Thumb-1 inhibitor; it was developed for HCV and approved in Japan in 2016 as part of a fixed-dose combination (19). Preclinical selectivity studies demonstrated that beclabuvir was inactive (EC_50_ >14 μM) against a diverse panel of non-HCV viruses, including poliovirus, rhinovirus, coronavirus, coxsackievirus, influenza, and HIV (20). During the COVID-19 pandemic, multiple groups used molecular docking and molecular dynamics simulations to predict that beclabuvir would inhibit SARS-CoV-2 RdRp (21–22). The compound demonstrated no antiviral activity in experimental assays (EC_50_ >20 μM) (23). This failure appeared to confirm the dogma that HCV NS5B Thumb-1 inhibitors could not have multi-family, broad-spectrum activity and that the Thumb-1 site was not a conserved, broad-spectrum antiviral target.

**Figure 1.**
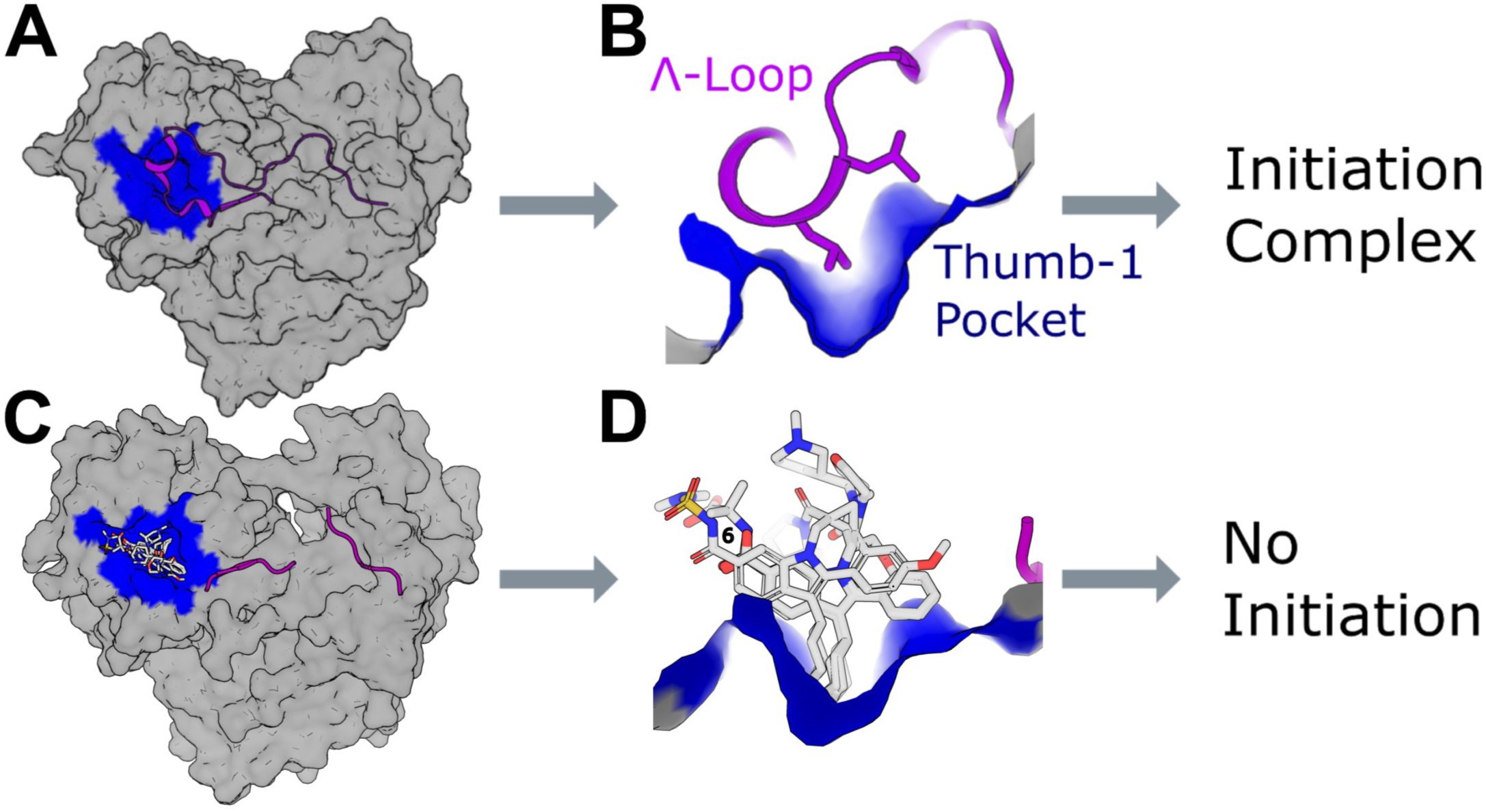
Thumb-1 Pocket Structure and Mechanism of Inhibition. **(A)** HCV NS5B apoprotein structure (PDB: 4AEP) shown as gray surface with Thumb-1 pocket (blue) and Λ1-loop (magenta). **(B)** Close-up of the apo Thumb-1 pocket showing endogenous leucine side chains from the Λ1-loop α-helix occupying the site. **(C)** Overlay of Thumb-1 inhibitors from co-crystal structures superimposed on holoprotein structure 2XWY. Seven of ten published Thumb-1 co-crystal ligands are shown (PDB: 2BRK, 2BRL, 2DXS, 2XWY, 3MWW, 4GMC, 4NLD); three structures (2WCX, 3Q0Z, 4DRU) are omitted for visual clarity. **(D)** Close-up of the Thumb-1 pocket with bound inhibitors demonstrating competitive displacement of the Λ1-loop. Inhibitor indole position C6 is indicated.

This study addresses whether the Thumb-1 pocket is confined to HCV NS5B or is a conserved and druggable broad-spectrum feature of the RNA-dependent RNA polymerase.

Here we report GALILEO’s discovery that the RdRp Thumb-1 allosteric pocket and its critical interaction with the Λ1-loop are structurally conserved across RNA viral families for the first time. We structurally validate this discovery using publicly available data from the Protein Data Bank (PDB), commercial software, and an open-source CE-align prompt that can be independently validated. Specifically, a fifteen amino acid Thumb-1 pocket was defined using ten HCV NS5B-ligand co-crystal structures available in the PDB. The pocket was then mapped onto an NS5B HCV apo structure in the PDB with complete homology. The PDB HCV NS5B apo structure was aligned with a PDB nsp12 SARS-CoV-2 apo structure using the annotations of the original authors of the PDB submissions. The alignment revealed structural conservation of the Thumb-1 pocket, the Λ1-loop and the Thumb-1/Λ1-loop interaction in HCV NS5B and SARS-CoV-2 nsp12. Simultaneously, we demonstrate that beclabuvir’s failure to demonstrate broad-spectrum activity reflects a pharmacophoric limitation of its indole C6 carbonyl substituent and the drug’s dependence on the HCV-specific residue R503, not a limitation of the Thumb-1 target itself.

We further demonstrate that broad-spectrum inhibition of the Thumb-1/Λ1-loop interaction is possible. To identify chemistry capable of cross-family Thumb-1 inhibition, we employed ChemPrint, an experimentally validated molecular-geometric deep learning model within the GALILEO drug discovery platform (24). The GALILEO platform and ChemPrint model used to discover MDL-001 have been peer-reviewed and published previously (25). Trained on a proprietary antiviral bioactivity dataset spanning multiple RNA viral families, ChemPrint identified novel Thumb-1 chemistry. MDL-001 does not contain an indole C6 carbonyl substituent, which Bristol Myers Squibb previously reported was required for NS5B engagement (20). MDL-001 inhibits both HCV and SARS-CoV-2. Mechanism-of-action studies confirm Thumb-1 engagement. The compound inhibits RNA synthesis in purified HCV and SARS-CoV-2 replication complexes (26) and selects for the canonical NS5B Thumb-1 resistance mutation in HCV replicons (27). Critically, this chemistry inhibits SARS-CoV-2 replication in vitro under conditions where beclabuvir remains inactive (23). These findings validate RdRp Thumb-1 as a druggable target and establish proof of concept for broad-spectrum non-nucleoside antiviral development.

This manuscript demonstrates MDL-001’s broad-spectrum antiviral activity against HCV and SARS-CoV-2. A previously published study validates these results in vivo. There, MDL-001 demonstrates multi-log antiviral efficacy against HCV, HBV, influenza, and SARS-CoV-2 in preclinical animal models and inhibits RSV and HDV in cell-based systems with nM EC90 potency (28). MDL-001 also demonstrates superiority or equivalence to multiple standards of care, including sofosbuvir, tenofovir, oseltamivir, remdesivir, nirmatrelvir, molnupiravir, and ribavirin in that study.

## 2. Results

### The Thumb-1 Pocket and Finger Loop Are Conserved across Viral Families

All findings reported in this section can be independently reproduced and validated in the public domain by any researcher from the given PDB entries, the alignment command, and the view given in the Methods.

#### Fifteen Residues Define a Consensus HCV NS5B Thumb-1 Pocket

We defined the residues that constitute the HCV Thumb-1 pocket using ten HCV NS5B/Thumb-1 inhibitor co-crystal structures found in the PDB.

Ten PDB structures of HCV NS5B bound to Thumb-1 inhibitors (**Supplemental Fig. S1A-C**) were analyzed. All NS5B residues within 5 Å of the bound ligands were identified (**Fig. 2**). Fifteen residues are within 5 Å in all published Thumb-1 co-crystal structures: ^392^LA^393^, ^395^AA^396^, T^399^, ^424^IL^425^, ^428^HF^429^, ^492^LGVP^495^, W^500^, and R^503^. The pocket is predominantly hydrophobic. Specifically, twelve of fifteen residues are hydrophobic ( ^392^LA^393^, ^395^AA^396^, ^424^IL^425^, F^429^, ^492^LGVP^495^, W^500^).

**Figure 2.**
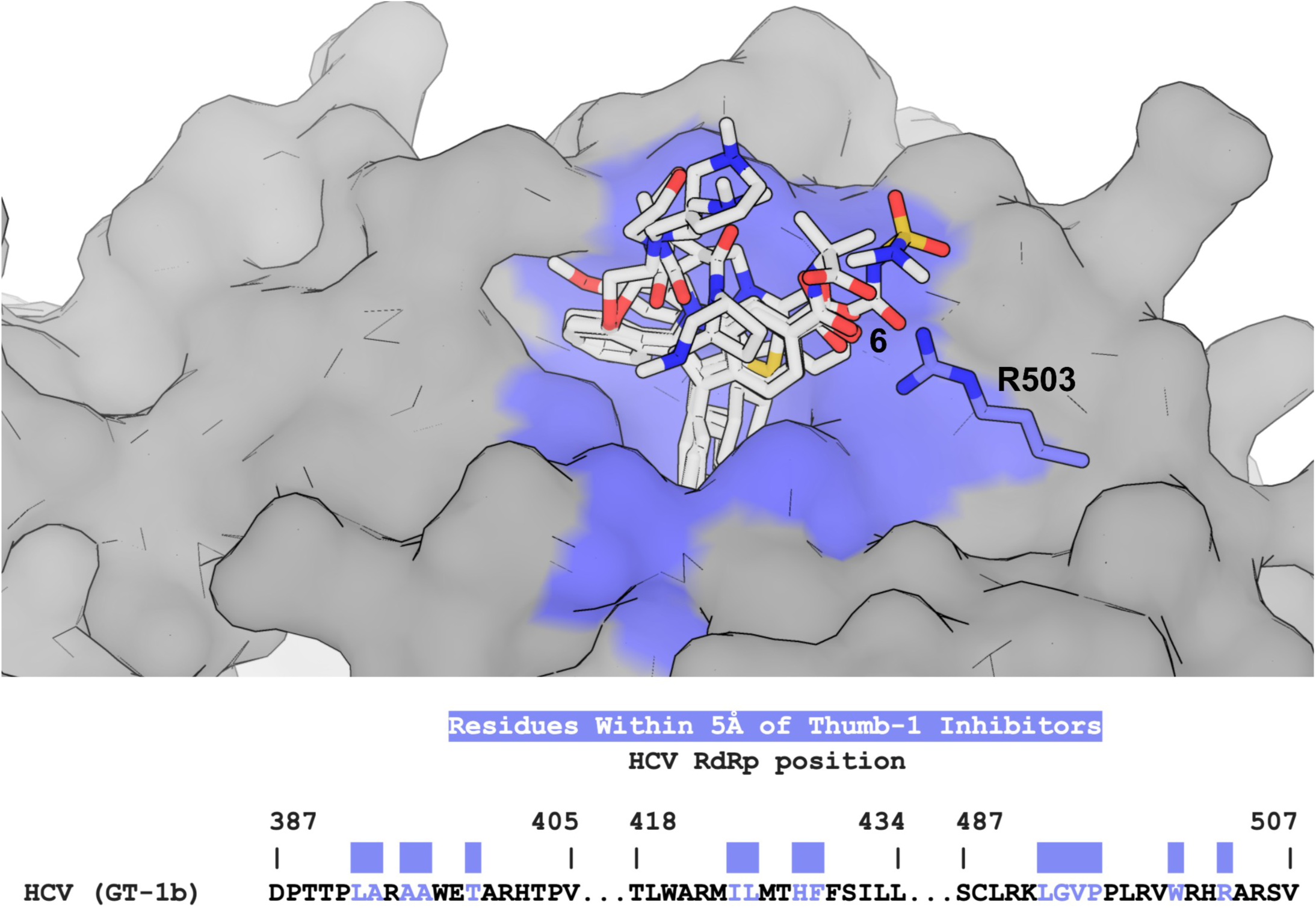
Co-Crystal Structure and Sequence of Inhibitor-Defined HCV NS5B Thumb-1 Pocket Residues. View of the 15 HCV NS5B residues within 5 Å (blue) of the bound ligand in the ten published Thumb-1 inhibitor co-crystal structures noted in Figure 1. The R^503^ side chain is shown as sticks. The 15 residues are also shown within the HCV Genotype 1b NS5B thumb domain sequence.

#### HCV NS5B Thumb-1 Pocket Residues are Conserved in SARS-CoV-2 nsp12

The fifteen amino acid HCV NS5B Thumb-1 pocket defined above was mapped onto an NS5B HCV apo structure (PDB 4AEP) with complete homology (**Fig. 3A**). The HCV NS5B apo structure was then aligned with a nsp12 SARS-CoV-2 apo structure (PDB 6M71) using the annotations of the original authors of the PDB submissions (29–30, 38). The alignment revealed structural-sequence conservation of the Thumb-1 pocket (**Fig. 3B-C**). SARS-CoV-2 nsp12 (PDB 6M71) and HCV NS5B (PDB 4AEP) present amino acids that occupy the same three-dimensional space (**Fig. 3D and 3E**). Specifically, ten of the fifteen HCV NS5B Thumb-1 pocket residues have a SARS-CoV-2 residue within 1.5 Å that is either identical or a like-for-like substitution (**Supplementary Table S1**) (39–42). Six of the ten pairs occupy the same position in the structure-based alignment (**Fig. 3D**). The remaining four pairs occupy the same three-dimensional position from different alignment columns (**Fig. 3E**). A sequence alignment cannot reveal these four pairs. Only the structure-based alignment reported here recovers them (**Fig. 3F**). This is the first study to demonstrate the structural conservation of the Thumb-1 pocket in a second viral family.

**Figure 3.**
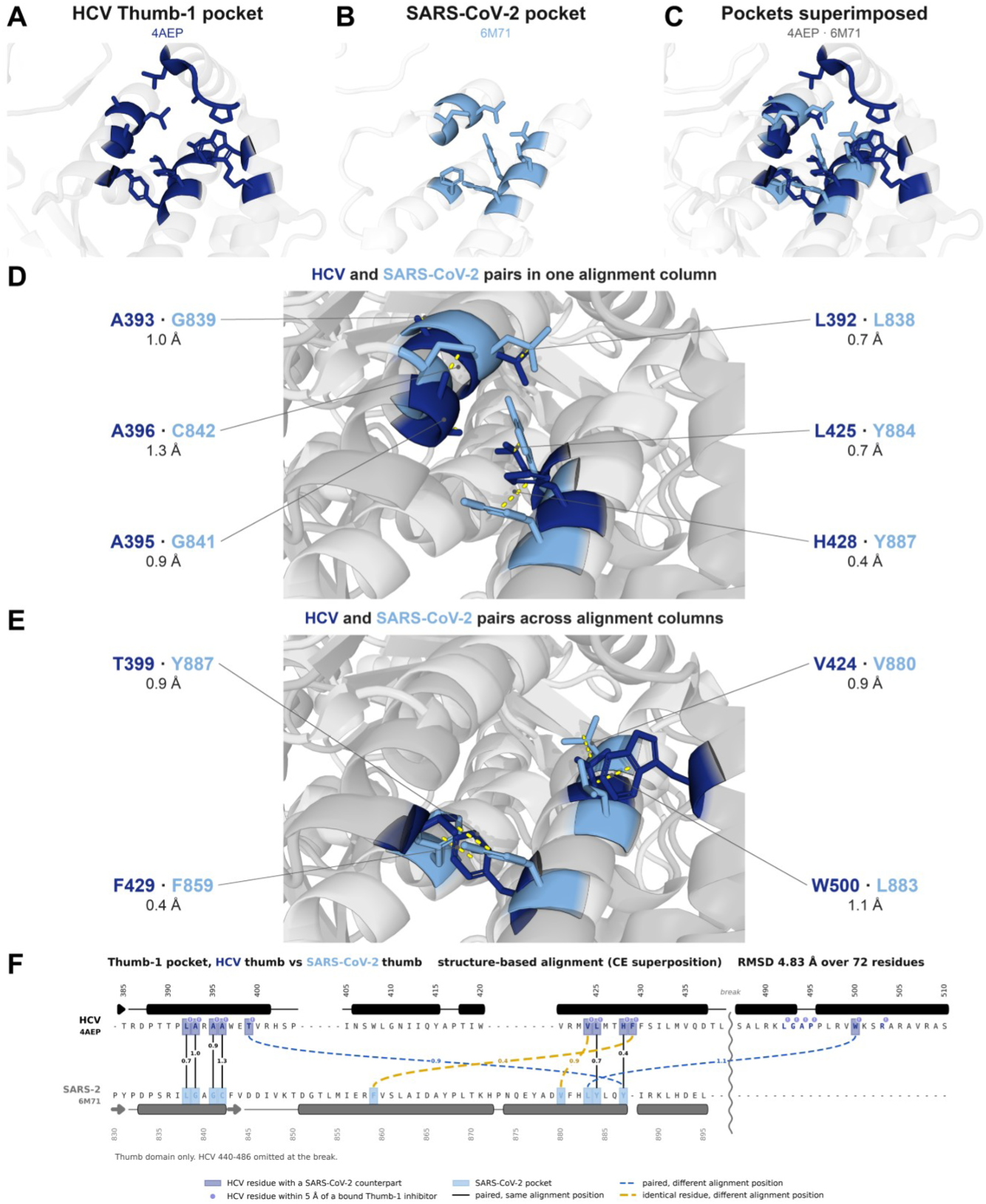
The Thumb-1 Pocket is Conserved Across Viral Families. The HCV NS5B thumb subdomain was assigned to residues 371 to 528 (38). The SARS-CoV-2 RdRp thumb subdomain was assigned to residues 816 to 920 (30). Here we align HCV NS5B PDB 4AEP to SARS-CoV-2 RdRp PDB 6M71 via the annotations from their respective authors to validate the GALILEO discovery. We defined the residues that constitute the HCV NS5B Thumb-1 pocket using ten HCV NS5B/Thumb-1 ligand co-crystal structures published in the PDB. All residues within 5 Å of the bound ligands were identified and defined the pocket. Every panel uses that one alignment view. HCV is dark blue and SARS-CoV-2 is light blue throughout both figures. The remaining backbone is gray. **(A)** The fifteen amino acids defined in the HCV NS5B co-crystals, mapped onto the HCV NS5B apo structure. **(B)** The SARS-CoV-2 nsp12 amino acids occupy the same three-dimensional space. **(C)** Both overlaid. **(D)** The six pairs sharing a structure-based alignment position. **(E)** The four pairs occupying the same three-dimensional position from a different position of that alignment. **(F)** Structure-based alignment across the pocket. The alignment used the full annotated thumb domains, HCV NS5B 371 to 528 against SARS-CoV-2 nsp12 816 to 920. This panel is cropped to the region that carries the Thumb-1 pocket, HCV NS5B 385 to 510 against SARS-CoV-2 nsp12 830 to 895. HCV 440 to 486 is omitted so the connectors fit. None of the fifteen pocket residues fall in the omitted stretch. A solid connector marks a shared alignment position. A blue dashed connector marks a pair drawn from differing alignment positions. An amber dashed connector marks an identical residue drawn from differing alignment positions. Each connector carries the distance between the two residues in Å. Shading marks an HCV NS5B Thumb-1 pocket residue that has a SARS-CoV-2 nsp12 counterpart. Two of the four pairs drawn from different positions carry the same amino acid: HCV F429 with SARS-CoV-2 F859, and HCV V424 with SARS-CoV-2 V880.

#### HCV NS5B Λ1-loop and its Thumb-1 Interface Are Conserved in SARS-CoV-2 nsp12

The Λ1-loop is the natural ligand for the Thumb-1 pocket in HCV NS5B. **Figure 4A** reports the HCV Λ1-loop in HCV NS5B PDB 4AEP, shown as the whole loop and, below it, as the tip helix seen down its axis and from two side views.

**Figure 4.**
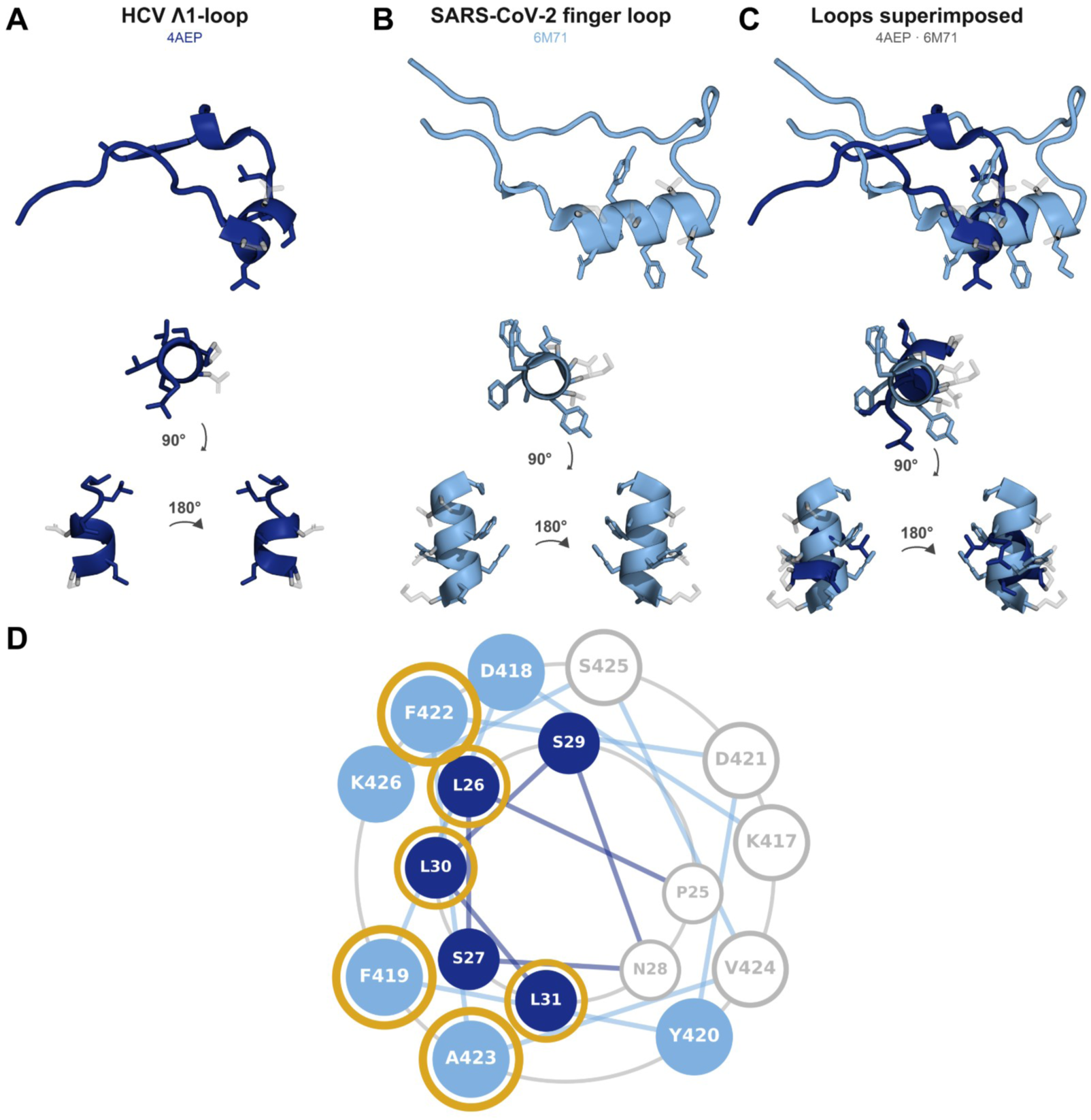
The Λ1-loop is Conserved Across Viral Families. The same superposition and the same alignment view as Fig. 3. The HCV Λ1-loop is dark blue and the SARS-CoV-2 finger loop is light blue. **(A)** The HCV Λ1-loop in PDB 4AEP, shown as the whole loop and, below it, as the tip helix down its axis and from two side views after the marked rotations. **(B)** The SARS-CoV-2 finger loop in PDB 6M71, in the same views. **(C)** The two loops superimposed, in the same views. **(D)** The two tips as overlaid wheels, HCV inner and SARS-CoV-2 outer, with pocket-facing positions filled and the rest open. Pocket-facing means any atom within 5 Å of the residue’s own pocket. A goldenrod ring marks a hydrophobic side chain that points into the pocket.

The Thumb domain alignment reported above identified a Finger Loop in SARS-CoV-2 PDB 6M71 (**Figure 4B**) without fitting. **Figure 4C** overlays the two structures. The centers of the two loop tips are 1.3 Å apart, and their nearest atoms are 0.3 Å apart. Each HCV tip Cα is within 4.0 Å of a SARS-CoV-2 tip Cα.

Each tip presents a hydrophobic face to the pocket (**Fig. 4D**). Each tip has four positions with a hydrophobic side chain. Three of the four amino acids point into the pocket in both PDB structures.

Taken together, the Thumb-1 pocket, the Finger Loop, and their interaction are conserved across HCV NS5B and SARS-CoV-2 nsp12. All reported findings above can be independently reproduced and validated in the public domain by any researcher from the given PDB entries, the alignment command, and the view given in the Methods.

### Identification of MDL-001 as a Broad-Spectrum Thumb-1 Inhibitor

The conservation of the Thumb-1 pocket in HCV and SARS-CoV-2 RdRps suggested the site could serve as a target for broad-spectrum antiviral inhibition across viral families. We leveraged GALILEO and ChemPrint to assemble a proprietary Thumb-1 training dataset and discover novel broad-spectrum inhibitor chemistry.

#### Discovery of a Broad-Spectrum Thumb-1 Data Set

We identified core scaffolds and leveraged them to construct a multi-feature pharmacophoric model of Thumb-1 inhibitors (**Fig. 5A**). The model was used to discover unknown Thumb-1 inhibitors and to construct a proprietary, broad-spectrum data set for ChemPrint model training (**Fig. 5A**).

**Figure 5.**
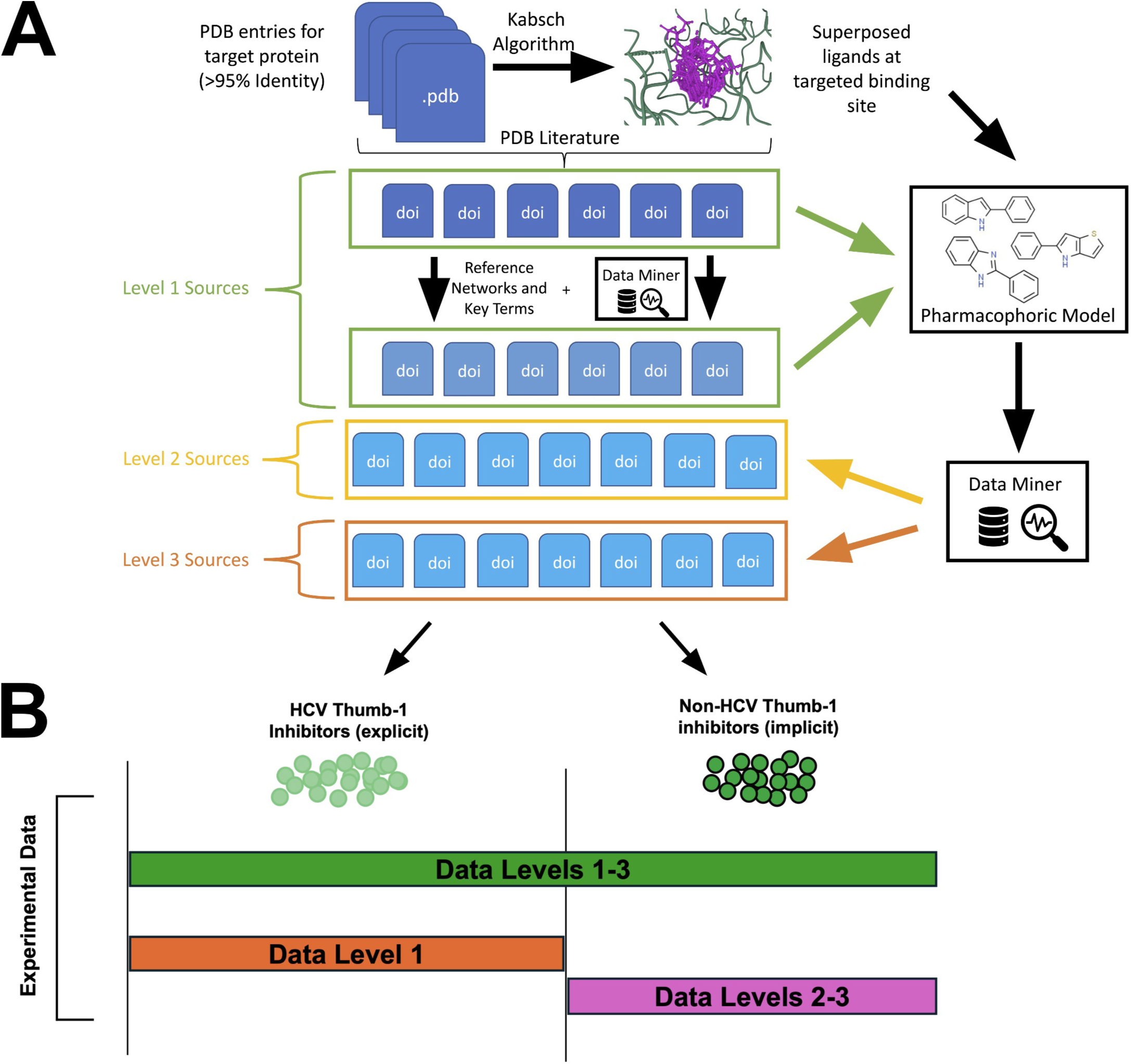
Pharmacophoric Model, Proprietary Data Discovery & ChemPrint Drug Discovery Training Datasets. **(A)** Multi-feature Thumb-1 inhibitor pharmacophoric model and proprietary data discovery **(B)** ChemPrint Drug Discovery Model Training Datasets.

Thumb-1 ligands in the PDB share one of three core scaffolds: 2-phenylindole, 2-phenylbenzimidazole, and 5-phenyl-4H-thieno[3,2-b]pyrrole. We leveraged these core scaffolds to construct a multi-feature pharmacophoric model and subsequently stratify the broader inhibitor landscape into a three-tiered classification hierarchy. Level 1 captured compounds from publications that explicitly reported Thumb-1 site inhibition in HCV NS5B. Level 2 captured molecules tested in cell-free RdRp assays for any virus whose structures matched the Thumb-1 pharmacophoric profile, but whose binding pocket was not explicitly identified. Level 3 captured molecules evaluated in cell-based infection-model assays against viruses bearing Thumb-1-associated scaffolds. Molecules discovered and placed in Level 2 and Level 3 datasets are proprietary to this discovery program. Together, these three levels created a proprietary broad-spectrum Thumb-1 ChemPrint training dataset (**Data Levels 1-3, Fig. 5B**) used to train ChemPrint. We designated Level 1 compounds as explicit Thumb-1 data (**Data Level 1, Fig. 5B**) and Level 2 and Level 3 compounds as implicit Thumb-1 data (**Data Levels 2-3, Fig. 5B**). Two additional ChemPrint models were trained to distinguish HCV-confined Thumb-1 chemistry from broad-spectrum Thumb-1 chemistry. The first model was trained on explicit Thumb-1 data (**Data Level 1, Fig. 5B**). The second model was trained on implicit Thumb-1 data (**Data Levels 2-3**, Fig. 5B).

#### Discovery of MDL-001 In Silico

ChemPrint trained on **Data Levels 1-3 (Fig. 5B)** was used to score broad-spectrum Thumb-1 activity on an inference library of clinically evaluated compounds and identify broad-spectrum inhibitors of the RdRp Thumb-1 site. ChemPrint’s prediction score for each inference compound represents the probability that a compound will demonstrate potent, broad-spectrum Thumb-1 activity. Top-scoring compounds were filtered for novelty to viral disease. MDL-001 (pipendoxifene; **Fig. 6A**) was predicted to have a broad-spectrum Thumb-1 mean inhibition probability of 0.76 ± 0.02 (95% CI). MDL-001 has established clinical safety and tolerability, oral bioavailability and novelty to antiviral applications (43). MDL-001 was nominated for additional *in silico* and *in vitro* validation based on these results.

**Figure 6.**
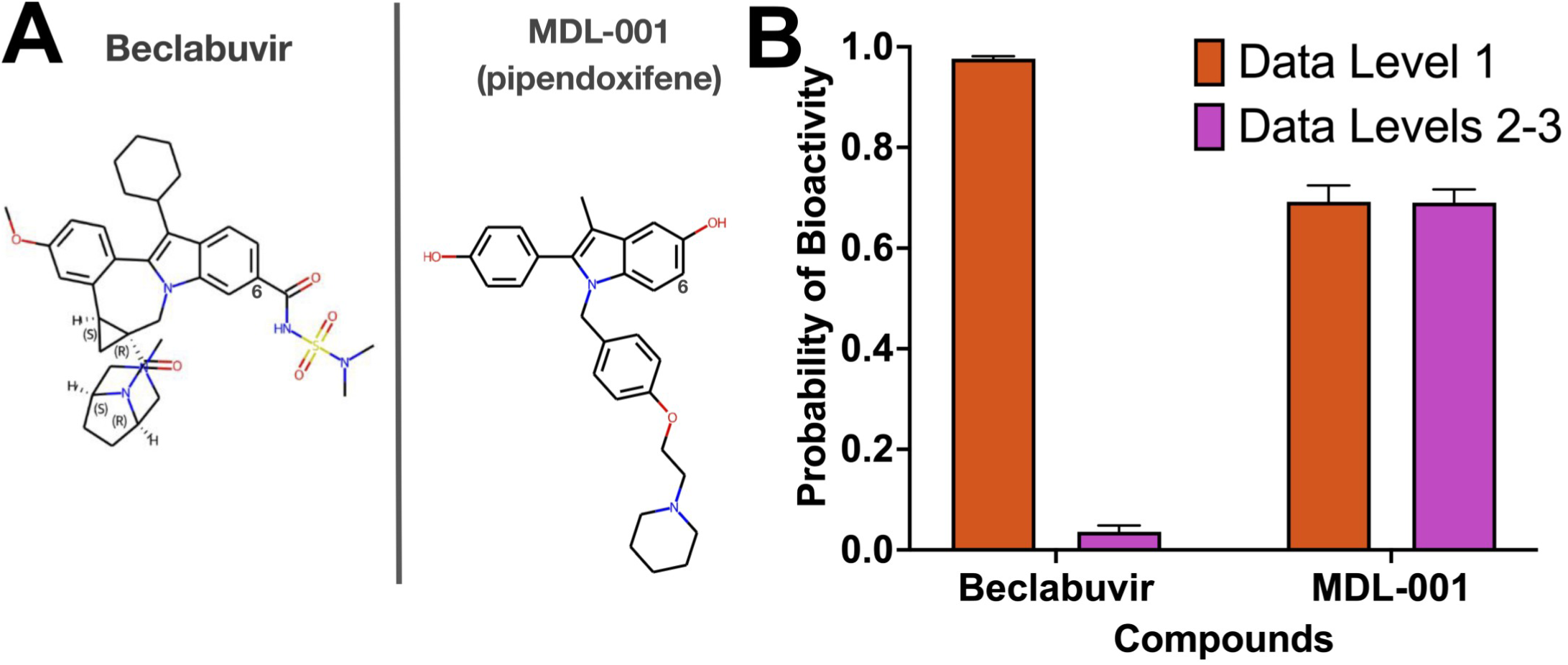
ChemPrint Prediction Scores Distinguish Broad-Spectrum Thumb-1 Inhibitors from HCV-Specific Thumb-1 Inhibitors. **(A)** Comparison of beclabuvir and MDL-001 chemical structures with indole C6 annotated. **(B)** Comparison of MDL-001 and beclabuvir prediction scores from models trained on explicit Thumb-1 data (**Data Level 1**) versus implicit Thumb-1 data (**Data Levels 2-3**). Error bars represent 95% confidence intervals. Beclabuvir scores 0.98 on Data Level 1 but only 0.04 on Data Levels 2-3, consistent with its lack of activity against SARS-CoV-2.

#### MDL-001 Broad-Spectrum Potential Relative to Beclabuvir

The approved Thumb-1 inhibitor beclabuvir and MDL-001 illustrate how ChemPrint models trained separately on Level 1 and Level 2-3 data distinguish broad-spectrum chemistry from HCV-restricted Thumb-1 chemistry (**Fig. 6**).

The chemistry and structure of beclabuvir and MDL-001 were compared to understand distinctions between the two compounds that may lead to distinct biological activity (**Fig. 6A**). Beclabuvir possesses an indole C6 carbonyl, consistent with all known indole-based HCV NS5B Thumb-1 inhibitors with micromolar or better activity against HCV (20, 44). MDL-001 lacks a C6 substituent entirely. Alternatively, it presents a hydroxyl at C5.

Models trained on explicit Thumb-1 data (**Data Level 1, Fig. 5B**) assigned high prediction scores of 0.69 and 0.98 to MDL-001 and beclabuvir, respectively (**Fig. 6B**). Similarly, molecular dynamics simulations indicated that both compounds have binding energies comparable to the native Λ1-loop (approximately -8.45 ± 0.65 kcal/mol), further supporting HCV Thumb-1 site engagement (**Supplemental Table S2**). The critical distinction emerged from models trained on implicit Thumb-1 data (**Data Levels 2-3, Fig. 5B**). These models learned chemical features associated with broad-spectrum Thumb-1 inhibition across non-HCV viral families and scored MDL-001 at 0.69, but beclabuvir at only 0.04 (**Fig. 6B**). This divergence indicates that MDL-001 carries chemical features compatible with Thumb-1 engagement across multiple viral families, whereas beclabuvir’s molecular features restrict its activity to HCV. The prediction from the implicit Thumb-1 model (**Data Levels 2-3, Fig. 5B**) for beclabuvir aligns with its experimental inactivity against SARS-CoV-2 (EC_50_ > 20 µM) (23) and literature-reported inactivity against poliovirus, rhinovirus, coxsackievirus, influenza virus, and HIV. This result validates that this model captures the chemistry responsible for cross-family Thumb-1 activity.

#### MDL-001 Represents a Novel Chemical Class Among RdRp and Thumb-1 Inhibitors

We analyzed MDL-001’s position in chemical space relative to all known RdRp inhibitors to determine whether MDL-001 constitutes a distinct chemical class. Principal Component Analysis (PCA) and t-distributed Stochastic Neighbor Embedding (t-SNE) projections of molecular embeddings together with pair-wise Tanimoto analysis revealed four findings (**Fig. 7A-B**).

**Figure 7.**
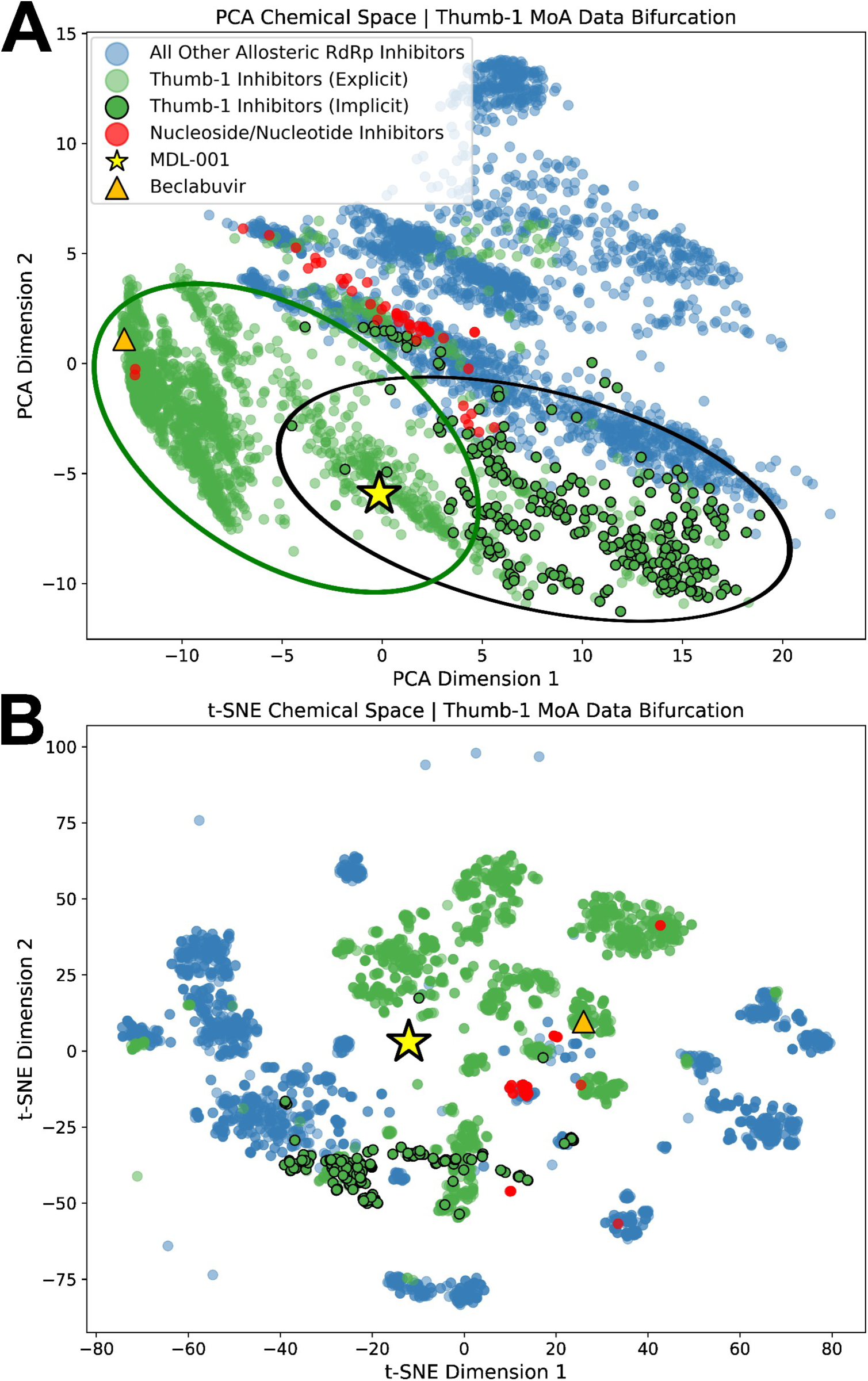
MDL-001 and Beclabuvir Occupy Distinct Regions of RdRp Inhibitor Chemical Space. **(A)** Principal component analysis (PCA) of molecular embeddings. Ovals delineate the trace of explicit HCV Thumb-1 chemical space (green) and implicit non-HCV Thumb-1 chemical space (black). **(B)** t-SNE projection of the same dataset.

Three observations emerged in PCA space, where Euclidean distance is preserved. First, MDL-001 occupies a region of chemical space distinct from RdRp inhibitors with non-Thumb-1 mechanisms of action. Second, MDL-001 occupies a region of PCA space between explicit HCV NS5B Thumb-1 inhibitors and implicit non-HCV Thumb-1 inhibitors, consistent with its predicted activity across viral families. Third, MDL-001 occupies a distant region of Thumb-1 chemical space from beclabuvir. Furthermore, the pair-wise Tanimoto similarity score between MDL-001 and beclabuvir is 0.15. This demonstrates that MDL-001 is chemically novel to beclabuvir and confirms that the two compounds reside in distinct regions of Thumb-1 chemical space. A fourth observation emerged in t-SNE space where local chemical neighborhood relationships are preserved. MDL-001 forms its own isolated cluster, indicating differentiated chemistry from all other plotted compounds. Together, this mathematically demonstrates that MDL-001’s chemistry is novel compared to all prior non-Thumb-1 RdRp ligand chemistry and both explicit HCV Thumb-1 ligand chemistry and implicit non-HCV Thumb-1 ligand chemistry, including beclabuvir.

### In vitro confirmation of MDL-001 Thumb-1 Mechanism of Action and Broad-Spectrum Activity

To determine whether MDL-001 inhibits viral replication through Thumb-1 engagement and to test its broad-spectrum potential, we performed biochemical and virological studies in both HCV and SARS-CoV-2.

#### MDL-001 Directly Inhibits HCV RNA Synthesis via NS5B

We evaluated MDL-001’s ability to directly inhibit viral replication activity using a purified HCV replication complex (RC) assay. The assay provides a stringent test of direct polymerase inhibition. The HCV RC assay is an established cell-free system that isolates functional replication machinery from cellular factors.

MDL-001 inhibited *de novo* RNA synthesis in purified RCs (**Fig. 8A**). This result demonstrates that MDL-001’s antiviral activity is a result of direct targeting of HCV NS5B (26).

**Figure 8.**
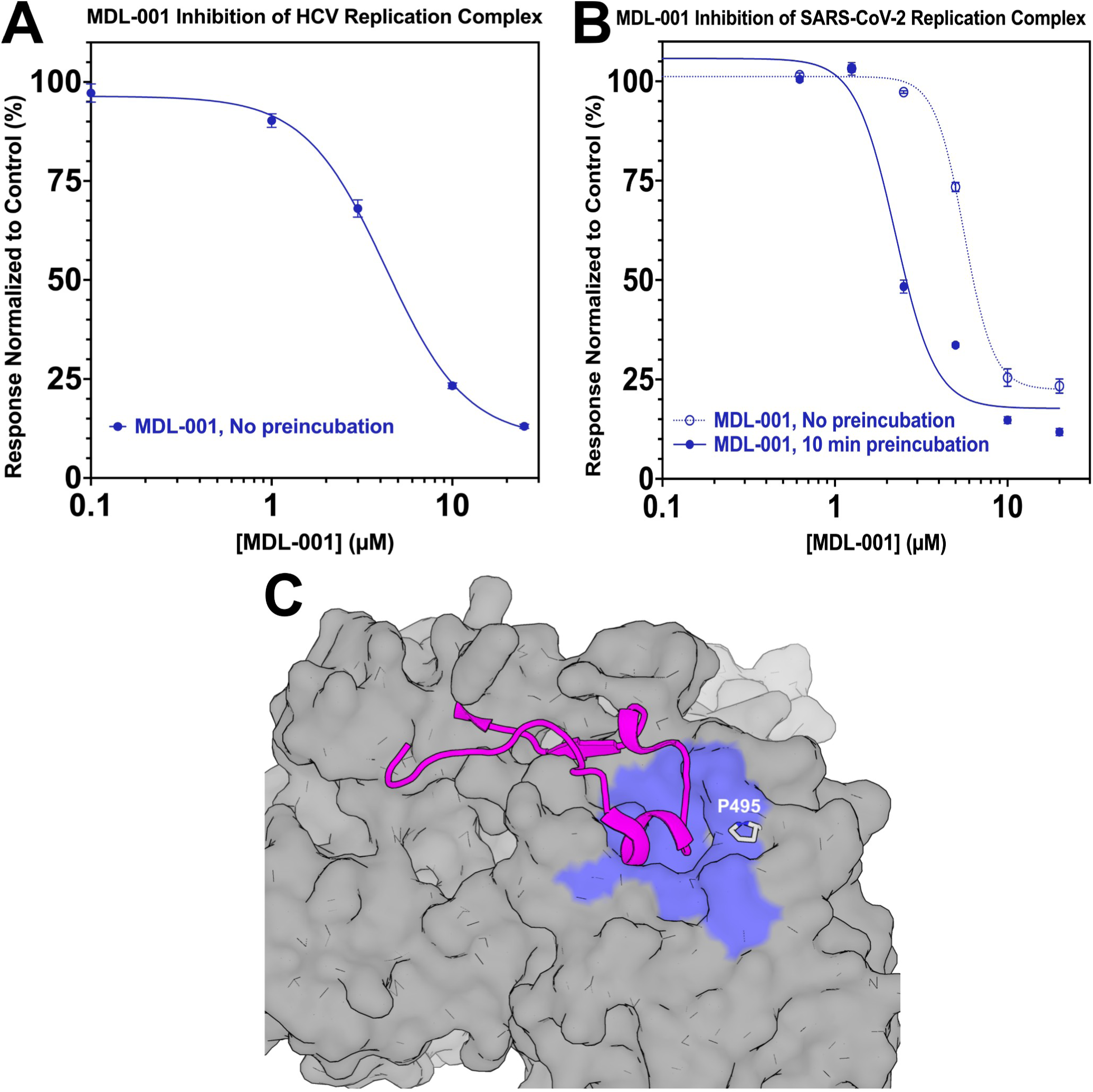
MDL-001 Directly Inhibits RNA Synthesis via RdRp in Purified HCV and SARS-CoV-2 Replication Complexes and Selects for the Canonical NS5B Thumb-1 Resistance Mutation, Confirming the Thumb-1 Mechanism of Action. **(A)** Inhibition of [α-³²P]CTP incorporation into nascent viral RNA by MDL-001 in purified HCV replication complexes. Data represent mean ± SD from three independent replicates. Omission of rNTPs abolished ³²P incorporation, confirming de novo RNA synthesis. The chain terminator 3’-dCTP reduced incorporation by >98% relative to DMSO, serving as a positive control for polymerase-dependent inhibition. **(B)** Inhibition of RNA synthesis by MDL-001 in purified SARS-CoV-2 replication complexes, measured after a 10-minute preincubation or with no preincubation before the reaction was started. Data represent mean ± SD from three independent replicates. (C) Location of NS5B residue P⁴⁹⁵ in the Thumb-1 allosteric pocket. Cartoon representation of HCV NS5B (PDB-ID: 2XXD; Λ1-loop = magenta, inhibitor-proximal residues = blue). P^495^ is represented as spheres at the rim of the inhibitor-binding pocket.

#### MDL-001 Directly Inhibits SARS-CoV-2 RNA Synthesis via nsp12

We evaluated MDL-001’s ability to directly inhibit broad-spectrum viral replication activity using a purified SARS-CoV-2 replication complex (RC) assay. MDL-001 inhibited *de novo* RNA synthesis in purified SARS-CoV-2 RCs (Fig. 8B). The observed potency of MDL-001 doubled with 10 minutes of pre-incubation. Beclabuvir produced no inhibition at concentrations up to 20 μM. This biochemical result demonstrates that MDL-001 directly inhibits the SARS-CoV-2 viral polymerase. The result also replicates the pre-incubation phenomenon reported for beclabuvir and HCV NS5B by Bristol Myers Squibb researchers (45). The implications are discussed below.

#### MDL-001 Selects for the Canonical Thumb-1 Resistance Mutation

We evaluated MDL-001’s ability to directly inhibit viral replication via the NS5B Thumb-1 pocket by mapping resistance mutations to the binding site.

Serial passage of HCV replicons under MDL-001 selective pressure yielded a convergent P495S substitution in three distinct colonies (Fig. 8C). Beclabuvir resistance selection in HCV replicons mapped to the same NS5B position 495 (P495A/S/L/T) (27). Substitutions at NS5B position 495 have previously been introduced into HCV replicons by site-directed mutagenesis. P495S raises the beclabuvir EC50 approximately 67-fold in a genotype 1a replicon (27). P495S is therefore an established, site-specific marker of Thumb-1 engagement.

P495S selection permitted replicon growth under MDL-001 concentrations 5 times its EC50 against WT HCV. Whole-genome sequencing of the resistant replicons identified no other convergent amino acid substitution across the three colonies, within or outside NS5B. All additional variants were non-coding or synonymous. This convergence establishes that both compounds engage the Thumb-1 site directly through a shared pocket interface.

#### MDL-001 Demonstrates Broad-Spectrum Antiviral Activity

We evaluated MDL-001’s broad-spectrum antiviral activity in vitro against HCV and SARS-CoV-2.

MDL-001 demonstrated an EC_50_ of 77 nM in an in vitro HCV replicon assay (**Table 1**). MDL-001 demonstrated an EC_50_ of 188 nM in an in vitro SARS-CoV-2 assay (**Table 1**). No cytotoxicity was observed across the antiviral concentration range. Beclabuvir demonstrated no activity up to 20,000 nM against SARS-CoV-2 (23). These results confirm MDL-001’s broad-spectrum activity and are consistent with ChemPrint predictions for both compounds.

**Table 1.** MDL-001 inhibits HCV and SARS-CoV-2 replication in cell-based antiviral assays. All values are the mean of three independent experiments (n = 3).

| Virus (strain) | Antiviral potency |  | Cytotoxicity |  |
| --- | --- | --- | --- | --- |
|  | EC<sub>50</sub> (nM) | EC<sub>90</sub> (nM) | CC<sub>50</sub> (nM) | CC<sub>10</sub> (nM) |
| HCV JFH-1 (genotype 2a) | 77 | 300 | >500 | >500 |
| SARS-CoV-2 (USA-WA1/2020) | 188 | 345 | >12,500 | 2,380 |

*HCV*: Huh-7.5.1 cells stably expressing the JFH-1 subgenomic luciferase reporter replicon (genotype 2a); replication was quantified by firefly luciferase activity and cell viability by CellTiter-Glo, each as a percentage of DMSO control. *SARS-CoV-2 (USA-WA1/2020)*: HeLa-ACE2 cells; antiviral activity was determined by immunostaining for viral nucleoprotein on a Celigo imaging cytometer with cytotoxicity assessed in parallel by MTT assay, each as a percentage of control.

## 3. Discussion

### The HCV NS5B Thumb-1 Pocket is a Conserved Allosteric Target Across Viral Families

RNA virus pandemics occur regularly and cause widespread death and economic devastation (1–4). No direct-acting, non-nucleoside antiviral has ever received regulatory approval for infections spanning multiple RNA viral families (1). The conventional view holds that allosteric sites on viral polymerases diverge too rapidly for cross-family targeting (13, 16–17). The failure of beclabuvir against poliovirus, rhinovirus, coronavirus, coxsackievirus, influenza, and HIV appeared to confirm this view (20). Our findings reported here overturn this assumption.

HCV NS5B and SARS-CoV-2 nsp12 present the same Thumb-1/Λ1-loop biology at the same points in three-dimensional space (**Fig. 2-4**). Ten of the fifteen HCV NS5B residues that define the HCV NS5B Thumb-1 pocket have a SARS-CoV-2 nsp12 counterpart within 1.5 Å (**Fig 3**). Six of the ten pairs occupy the same position in the structure-based alignment. The remaining four pairs occupy the same three-dimensional position from different alignment columns. This result suggests convergent evolution of the Thumb-1 pocket across otherwise divergent viral families. Thumb domain alignment and superposition of HCV NS5B and SARS-CoV-2 nsp12 PDB structures demonstrate the presence of a Finger Loop in each protein as well (**Fig. 4**). Both Finger Loops originate outside the thumb domain region and superimpose onto each other under Thumb domain alignment. Each loop reaches across the protein and ends in a short helix. Both present a hydrophobic face to the pocket. Together with the pocket residue conservation reported above, this suggests convergent evolution of the functional biology of the Thumb-1/Λ1-loop interaction.

The proprietary GALILEO AI platform discovered that the broad-spectrum RdRp Thumb-1 allosteric pocket and its critical interaction with the Λ1-loop are structurally conserved across RNA viral families. However, the validation of the discovery reported here was executed using publicly available data from the Protein Data Bank (PDB), commercial software, and an open-source CE-align prompt. Thus, all HCV NS5B/ SARS-CoV-2 nsp12 Thumb-1 findings reported here can be independently recreated and validated in the public domain by any researcher. We have provided a link in the Methods that automates this process.

### MDL-001 Represents a Novel Chemical Class whose Divergence from Known Thumb-1 Pharmacophores is Required for Broad-Spectrum Activity

The failure of first-generation Thumb-1 chemistry outside HCV NS5B reflects a pharmacophoric limitation rather than a target limitation. R^503^ anchors all previously known indole-based Thumb-1 inhibitors through a hydrogen bond with their C6 carbonyl (**Fig. 2 & 6A)** (20, 44). R^503^ is absent from coronaviruses (**Fig. 3F**). Beclabuvir’s inactivity against SARS-CoV-2 and a broad panel of non-HCV viruses (20, 23) is explained by its dependence on the R ^503^ -carbonyl interaction for antiviral activity. MDL-001 lacks a C6 substituent and therefore does not rely on a C6 carbonyl-R^503^ interaction for Thumb-1 engagement (**Fig 6A**).

Chemical space analysis confirms this novelty. MDL-001 and beclabuvir share a Tanimoto similarity of only 0.15 and occupy distinct regions in PCA and t-SNE projections of molecular embeddings (**Fig. 7A-B**). ChemPrint models trained on implicit Thumb-1 data from non-HCV viral families scored MDL-001 at 0.69, but beclabuvir at only 0.04 (**Fig. 6B**). This divergence confirms that MDL-001 carries chemical features compatible with cross-family Thumb-1 engagement, while beclabuvir’s molecular features restrict its activity to HCV.

### MDL-001 Inhibits Viral Replication through the Novel Broad-Spectrum Thumb-1 Pocket

MDL-001 inhibited de novo RNA synthesis in purified HCV replication complexes (**Fig. 8A**). This cell-free system assays functional replication machinery isolated from replicon cells and confirms direct polymerase targeting (26). MDL-001 also inhibited RNA synthesis in purified SARS-CoV-2 replication complexes. Preincubation of 10 minutes was observed to double MDL-001’s observed potency (Fig. 8B). Preincubation dependence is the kinetic signature of Thumb-1 inhibition and was the basis on which the mechanism was established for the HCV polymerase (45–46). This is the first biochemical demonstration of Thumb-1 inhibition outside HCV.

Serial passage of HCV replicons under MDL-001 selective pressure yielded the P495S substitution in three distinct colonies (Fig. 8C). P495S is the canonical Thumb-1 resistance mutation identified for beclabuvir and other Thumb-1 inhibitors (27). P495S selection permitted replicon growth under sustained MDL-001 pressure 5-fold above the EC50 concentration. MDL-001 and beclabuvir share a Tanimoto similarity of only 0.15 yet converge on the same resistance determinant at the same pocket residue (27, 45). This convergence establishes that MDL-001 engages the Thumb-1 site directly.

MDL-001 inhibited in vitro HCV replication with an EC_50_ of 77 nM (**Table 1**) and in vitro SARS-CoV-2 replication with an EC_50_ of 188 nM (**Table 1**). Beclabuvir was inactive up to 20,000 nM against SARS-CoV-2 (23). These results validate both the target discovery and the chemical design rationale.

### Conclusions: The RdRp Thumb-1 Pocket is a Conserved Target for Broad-Spectrum Antiviral Development

The development of a direct-acting broad-spectrum non-nucleoside inhibitor has been considered impossible (16–17). Conventional wisdom holds that allosteric sites on viral polymerases are poorly conserved, restricting non-nucleoside inhibitors to genotype- or species-specific activity (13).

Here we report GALILEO’s discovery that the RdRp Thumb-1 allosteric pocket and its critical interaction with the Λ1-loop are structurally conserved across RNA viral families for the first time. We further demonstrate that broad-spectrum inhibition of the Thumb-1/Λ1-loop interaction is possible with MDL-001.

MDL-001 studies reported elsewhere have demonstrated multi-log antiviral efficacy against HCV, HBV, influenza, and SARS-CoV-2 in preclinical animal models and inhibition of RSV and HDV in cell-based systems with nM EC_90_ potency (28). MDL-001 has also demonstrated in vivo superiority or equivalency to multiple standards-of-care including sofosbuvir, tenofovir oseltamivir, remdesivir, nirmatrelvir, and molnupiravir. These results confirm that the Thumb-1 mechanism identified here translates from purified replication complexes and in vitro assays to whole-organism infection across multiple viral families.

The conserved target family identified here provides the biological foundation for a Thumb-1 New Chemical Entity (NCE) discovery program (47). Successive, multi-scaffold chemical series have already been discovered by GALILEO/ChemPrint, reduced to practice, and validated in vitro against multiple viruses.

These findings establish RdRp Thumb-1 as a conserved allosteric pocket and a druggable target for broad-spectrum antiviral development for the first time.

## 4. Materials and Methods

### Structural Analysis of the Thumb-1/Λ1-loop Interface

Structural visualization and analysis were performed using PyMOL (Figs. 1-4, 8C). The HCV RdRp apoprotein structure (PDB: 4AEP) was rendered as a gray surface with the Thumb-1 pocket colored blue and the Λ1-loop colored magenta (**Fig. 1A-B**). Ten co-crystal structures of HCV NS5B bound to Thumb-1 inhibitors (PDB: 2BRK, 2BRL, 2DXS, 2XWY, 2WCX, 3MWW, 3Q0Z, 4DRU, 4GMC, 4NLD) were superimposed on holoprotein structure 2XWY to visualize inhibitor binding and Λ1-loop displacement (**Fig. 1C-D**). Residues within 5 Å of all ten co-crystal ligands were rendered to define the consensus Thumb-1 pocket (**Fig. 2**).

### Structural Alignment Method

Coordinates were taken from HCV NS5B (PDB: 4AEP)(29) and SARS-CoV-2 nsp12 (PDB: 6M71) (30). Chain A was kept in each case. Waters, hydrogens and alternate conformers were removed. The thumb domains were superimposed with the combinatorial extension algorithm (31) as implemented in PyMOL cealign (CE). The algorithm aligns on Cα atoms. SARS-CoV-2 was held fixed and HCV was moved onto it.

The alignment view is available at https://huggingface.co/spaces/Model-Medicines/rdrp-thumb-1.

PyMOL implementation was executed as follows:

fetch 4AEP, hcv
fetch 6M71, cov
remove hcv and not (polymer and chain A)
remove cov and not (polymer and chain A)
cealign cov and resi 816-920, hcv and resi 371-528

The superposition in **Figure 3C** was viewed as follows:

set_color hcv_blue, [0.106, 0.184, 0.541]
set_color cov_blue, [0.498, 0.690, 0.878] bg_color white
hide everything
show cartoon, hcv and resi 371-528
show cartoon, cov and resi 816-920 color gray80, hcv
color gray85, cov
set cartoon_transparency, 0.83
select hcv_pocket, hcv and resi 392+393+395+396+399+424+425+428+429+492+493+494+495+500+503 select cov_pocket, cov and resi 838+839+841+842+859+880+883+884+887
set cartoon_transparency, 0, hcv_pocket or cov_pocket
show sticks, (hcv_pocket or cov_pocket) and (sidechain or name CA)
color hcv_blue, hcv_pocket
color cov_blue, cov_pocket
set cartoon_side_chain_helper, 1
set stick_radius, 0.28 set_view (\
0.0079710, 0.9782301, 0.2073704,\
- 0.9678253, 0.0597035, -0.2444380,\
- 0.2514974, -0.1987498, 0.9472317,\
0.0000000, 0.0000000, -86.5812149,\
107.8475723, 94.2121048, 143.5296478,\
68.2612915, 104.9011383, -20.0000000)
fetch 6M71, cov
remove hcv and not (polymer and chain A)
remove cov and not (polymer and chain A)
cealign cov and resi 816-920, hcv and resi 371-528

The same camera (set view) vector was used to show both the pocket alignment and the loop alignment across **Figures 3** and **4**.

### ChemPrint Model Training and Compound Selection

NS5B/Thumb-1 inhibitor co-crystal structures were retrieved from the PDB. Bound ligands were superposed at the Thumb-1 binding site using the Kabsch algorithm to extract shared pharmacophoric features and construct a multi-feature Thumb-1 inhibitor pharmacophoric model (**Fig. 5A**). This model was deployed with reference network analysis and automated data mining to identify additional and unlabeled Thumb-1-associated compounds in the published literature and construct the proprietary training dataset described below.

Bioactivity data for Thumb-1 inhibitors were curated at three levels (**Fig 5B**): explicit HCV Thumb-1 data, RdRp inhibitors matching Thumb-1 pharmacophoric profiles (2-phenylindole, 2-phenylbenzimidazole, 5-phenyl-4H-thieno[3,2-b]pyrrole), and antiviral screening hits bearing Thumb-1-associated scaffolds. Compounds with bioactivity below 1 μM were classified as active. ChemPrint, a graph convolutional network for molecular activity prediction, was trained and validated as previously described (24).

Three models were trained on data subsets to assess differential predictive behavior (**Fig. 6**). The combined model (**Data Levels 1-3**) achieved an AUROC of 0.93. The explicit Thumb-1 model (**Data Level 1**) achieved an AUROC of 0.93. The implicit Thumb-1 model (**Data Levels 2-3**) achieved an AUROC of 0.76; the smaller dataset required an 85-15% random split rather than the t-SNE-based split used for the other models. The trained model screened approximately 17,000 compounds with Phase I clinical data.

### Chemical Space Analysis

Molecular embeddings for all RdRp inhibitors were generated using ChemPrint’s learned representations (24). Dimensionality reduction was performed using PCA and t-SNE (**Fig. 7**). PCA was computed on the full embedding matrix; the first two principal components explained 24% of the total variance. t-SNE projections were generated to visualize local neighborhood structure. Tanimoto similarity between MDL-001 and beclabuvir was calculated using Morgan fingerprints (radius 2, 2048 bits). Compounds were classified as explicit Thumb-1 inhibitors (**Data Level 1**), implicit Thumb-1 inhibitors (**Data Levels 2-3**), other allosteric RdRp inhibitors, or nucleoside/nucleotide inhibitors.

### Molecular Dynamics Simulations

MD simulations were performed on HCV RdRp (PDB: 2XHU) using AMBER with GPU-accelerated generalized Born implicit solvation (32). The structure was protonated using the H++ server (33) and prepared with tLEaP. Simulations comprised minimization, heating (10 ns), and production (100 ns) in triplicate. Binding energies for the Λ1-loop, beclabuvir, and MDL-001 were calculated using MMPBSA (34). (**Supplemental Table S2**).

### Purified HCV Replication Complex Assay

Membrane-associated replication complexes were isolated from HCV replicon cells as previously described (26, 35) and assayed by the in vitro replicase method of Chatterji and colleagues (36). Replication complex activity was measured by incorporation of [α-³²P]CTP into nascent RNA (**Fig. 8A**). Reactions contained 20 mM Tris-HCl pH 8, 10 mM MgCl₂, 5 mM DTT, 5 mM KCl, 40 μg/mL actinomycin D, 16 mM creatine phosphate, 40 ng/μL creatine kinase, 1 U RNasin, 1 mM each ATP and UTP, 5 mM GTP, 10 μM CTP, and 20 μCi [α-³²P]CTP. A 30 μL replication complex input was tested in triplicate. Two negative controls were included: omission of rNTPs to confirm that ³²P incorporation reflects de novo RNA synthesis, and 3’-dCTP (10 μM) as a chain terminator positive control for polymerase-dependent inhibition. DMSO served as the vehicle control. Following incubation at 35°C for 60 minutes, RNA was extracted and analyzed by denaturing gel electrophoresis and phosphorimaging. Responses were normalized to the DMSO control and fitted by four-parameter logistic regression with the top and bottom plateaus unconstrained.

### Purified SARS-CoV-2 Replication Complex Assay

Replication complexes were purified from SARS-CoV-2 (USA-WA1/2020) grown in B7 cells, a single-cell clone of ACE2-expressing HEK293T cells carrying a protease-activatable luminescent reporter (37), at a multiplicity of infection of 10 with harvest 48 hours after infection by the procedure used for the HCV replication complex above (35–36), adapted to SARS-CoV-2. Replication complex activity was measured by incorporation of [α-32P]CTP into nascent RNA (Fig. 8B). Reactions contained 20 mM Tris-HCl pH 8, 10 mM MgCl2, 5 mM DTT, 5 mM KCl, 40 μg/mL actinomycin D, 16 mM creatine phosphate, 40 ng/μL creatine kinase, 1 U RNasin, 1 mM each ATP and UTP, 5 mM GTP, 10 μM CTP, 20 μCi [α-³²P]CTP and a 30 μL replication complex input in a 50 μL reaction held at 35°C for 60 minutes. Each condition was tested in triplicate. MDL-001 and beclabuvir were titrated from 0.625 to 20 μM. Beclabuvir produced no inhibition at any concentration tested, with or without preincubation. In the preincubation condition, compound was combined with the replication complexes and held for 10 minutes at 35°C before RNA synthesis was initiated. In the no-preincubation condition, compound was added to the reaction mixture immediately before addition of the replication complexes. Remdesivir (10 μM) was the positive control and reduced incorporation to near-background levels. Two negative controls were included: omission of rNTPs to confirm that ³²P incorporation reflects de novo RNA synthesis, and 3’-dCTP (10 μM) as a chain terminator positive control for polymerase-dependent inhibition. DMSO served as the vehicle control. Radiolabeled product was purified and quantified by autoradiography. Responses were normalized to the DMSO control within each preincubation condition and IC50 values were determined by four-parameter logistic regression with the top and bottom plateaus unconstrained.

### Selection of MDL-001-Resistant HCV Replicons

Huh-7.5.1 human hepatoma cells stably expressing the JFH-1 subgenomic replicon (HCV genotype 2a) were maintained under the culture conditions described in the HCV Assay section below. Cells were seeded and cultured with increasing concentrations of MDL-001, starting at 0.5x EC_50_. Drug concentration was increased stepwise over 8 weeks until cells grew stably in the presence of MDL-001 at a concentration of 5x starting EC _50_. Resistant replicon RNA was extracted, and the NS5B RdRp coding region was amplified by RT-PCR. Amplicons were sequenced to identify resistance-associated substitutions (Fig. 8C).

### HCV Antiviral Assay

Huh-7.5.1 human hepatoma cells stably expressing the JFH-1 subgenomic luciferase reporter replicon (HCV genotype 2a) were maintained in Dulbecco’s Modified Eagle Medium (DMEM) supplemented with 10% fetal bovine serum (FBS), nonessential amino acids, 2 mM L-glutamine, 10 mM HEPES, 100 units/mL penicillin, 100 μg/mL streptomycin, and 300 μg/mL G418 at 37°C in a humidified 5% CO_2_ atmosphere. Cells were seeded in 96-well plates and treated with serial dilutions of MDL-001 for 72 hours. Triplicate wells were used for each compound concentration. DMSO-treated wells served as the vehicle control for normalization. HCV replication was quantified by measuring firefly luciferase activity in cell lysates and expressed as a percentage of the DMSO control. Cell viability was assessed in a separate experiment, with uninfected wells treated with identical compound dilutions, using the CellTiter-Glo luminescent cell viability reagent (**Table 1**).

### SARS-CoV-2 Antiviral Assay

Antiviral activity against SARS-CoV-2 was determined by immunostaining for viral nucleoprotein. HeLa-ACE2 cells were seeded at 2,000 cells per well in 96-well plates in DMEM supplemented with 10% FBS and incubated for 24 hours at 37°C with 5% CO2. Two hours before infection, the medium was replaced with 100 μL of DMEM supplemented with 2% FBS containing MDL-001 at six concentrations, with a DMSO control. SARS-CoV-2 (USA-WA1/2020, BEI Resources NR-52281) was added at 1,000 PFU in 50 μL of DMEM supplemented with 2% FBS, a multiplicity of infection of 0.25. Viral stocks were propagated in Vero-TMPRSS2 cells, sequence-verified, and titered before use. Plates were incubated for 48 hours at 37°C. Supernatants were removed and cells were fixed with 4% formaldehyde. Cells were immunostained for the viral nucleoprotein with a DAPI counterstain. Infected cells and total cells were quantified on a Celigo imaging cytometer. Percent infection was calculated as infected cells divided by total cells, background subtracted, with the DMSO control set to 100% infection. Cytotoxicity was measured concurrently in uninfected HeLa-ACE2 cells at the same compound dilutions by MTT assay. Curves were fit by nonlinear regression in GraphPad Prism. Nirmatrelvir and DMSO controls were included in every experiment. All assays were performed in biologically independent triplicates (**Table 1**).

### FBS Protein Binding Determination

Protein binding of MDL-001 was determined by rapid equilibrium dialysis (RED) in fetal bovine serum (FBS). MDL-001 (1 μM) was incubated with 2% and 10% FBS under both heat-inactivated and non-heat-inactivated conditions. The unbound fraction (F_ub_) at 10% FBS was 5.0% (heat-inactivated) and 4.7% (non-heat-inactivated). The unbound fraction at 2% FBS was 25.0% (heat-inactivated) and 24.4% (non-heat-inactivated). Mean recovery ranged from 91.2% to 112.4% across all conditions. These values were used to calculate free-fraction antiviral potency from the cell culture antiviral EC_50_ values reported above.

## Acknowledgments

The authors thank Dr. George Nicola, co-inventor with Daniel Haders, of MDL-001. The authors thank Dr. David A. Poole III and Dr. Daan P. Geerke for performing the molecular dynamics simulations at Vrije Universiteit, Dr. Kris White for performing *in vitro* SARS-CoV-2 antiviral testing at Icahn School of Medicine at Mount Sinai Department of Microbiology and Dr. Michael Bobardt for performing *in vitro* HCV assays, and mechanism studies with replication complex and resistant mutations at Scripps Research Institute. We also extend our gratitude to Dr. Tushar Menon, Dr. Launa Aspeslet, and Dr. David Garvey for their expert consultation and detailed review of this manuscript.

## Data Availability Statement

The Level 1 Training Set References are available in the Supplemental Spreadsheet. The alignment view is available at https://huggingface.co/spaces/Model-Medicines/rdrp-thumb-1.

