## Supplementary Material for "The RdRp Thumb-1 Pocket is a Conserved Target for Broad-Spectrum Antiviral Development"

### Supplementary Tables

#### *Supplementary Table S1. Residue Pair Hydrophobicity*

| Residue Pairs<br>(HCV * SARS-CoV-2) | Residue Pair Hydrophobicity: | Comments: |
|---|---|---|
| L392 * L838 | Identical | Identical |
| V424 * V880 | Identical | Identical |
| F429 * F859 | Identical | Identical |
| A393 * G839<br>A395 * G841 | Rose et al. (30):<br>● A = 0.74<br>● G = 0.72<br><br>Janin (31):<br>● A = 1.7<br>● G = 1.8 | Residues are immediately adjacent on the Janin and Rose scales. Janin reports that residues have identical transfer free energies of +0.3 kcal/mol. |
| A396 * C842 | Kyte-Doolittle (32):<br>● A = +1.8<br>● C = +2.5 | The Kyte-Doolittle scale classifies both residues as hydrophobic. |
| H428 * Y887 | Rose et al. (30):<br>● H = 0.78<br>● Y = 0.76<br><br>Janin (31):<br>● H = 0.8<br>● Y = 0.5 | Residues are immediately adjacent on the Rose scale. Both are classified as neutral on the Rose and Janin scales. |
| T399 * Y887 | Rose et al. (30):<br>● T = 0.70<br>● Y = 0.76<br><br>Janin (31):<br>● T = 0.7<br>● Y = 0.5 | Both residues are classified as neutral on the Rose and Janin scales. |
| W500 * L883 | Rose et al. (30):<br>● W = 0.85<br>● L = 0.85<br><br>Janin (31):<br>● W = 1.6<br>● L = 2.4 | Rose scale values are identical. Residues are classified as hydrophobic on both the Rose and Janin scales. |
| L425 * Y884 | Casari-Sippl SDRH (33)<br>● L = 0.5<br>● Y = 0.5 | Both residues have identical scores on the Casari-Sippl scale. |

***Supplementary Table S2.*** *Molecular mechanics combined with the Poisson-Boltzmann Surface Area (MMPBSA) for WT NS5B against the native $\Lambda$1-Loop, Beclabuvir, and MDL-001.*

| **Thumb-1 Binding Partner** | $\Delta E_{bind}$ **WT (kcal mol$^{-1}$)** |
|---|---|
| $\Lambda$1-Loop | -8.21 $\pm$ 0.16 |
| Beclabuvir | -8.33 $\pm$ 0.68 |
| MDL-001 | -8.45 $\pm$ 0.65 |

#### Supplementary Figures

**A**

##### Ligand Proximity to Thumb-1 Pocket: Amino Acids 361-420

| | | |
|---|---|---|
| PDB: 2BRK CMF | ELITSCSSNVSVAHDASGKRVYYLTRDPTTPLARAAWETARHTPVNSWLGNIIMYAPTLW | 420 |
| PDB: 2BRL POO | ELITSCSSNVSVAHDASGKRVYYLTRDPTTPLARAAWETARHTPVNSWLGNIIMYAPTLW | 420 |
| PDB: 2DXS JTP | ELITSCSSNVSVAHDASGKRVYYLTRDPTTPLARAAWETARHTPVNSWLGNIIMYAPTLW | 420 |
| PDB: 2XWY IB8 | ELITSCSSNVSVAHDASGKRVYYLTRDPTTPLARAAWETARHTPVNSWLGNIIMYAPTLW | 420 |
| PDB: 2WCX VGC | ELITSCSSNVSVAHDASGKRVYYLTRDPTTPLARAAWETARHTPVNSWLGNIIMYAPTLW | 420 |
| PDB: 3MWW BIW | ELITSCSSNVSVAHDASGKRVYYLTRDPTTPLARAAWETARHTPINSWLGNIIMYAPTLW | 420 |
| PDB: 4DRU 0LN | ELITSCSSNVSVAHDASGKRVYYLTRDPTTPLARAAWETARHTPINSWLGNIIMYAPTLW | 420 |
| PDB: 4GMC 1BI | ELITSCSSNVSVAHDASGKRVYYLTRDPTTPLARAAWETARHTPINSWLGNIIMYAPTLW | 420 |
| PDB: 4NLD 2N7 | ELITSCSSNVSVAHDASGKRVYYLTRDPTTPLARAAWETARHTPVNSWLGNIIMYAPTLW | 420 |
| PDB: 5TRK 23E | ELITSCSSNVSVAHDASGKRVYYLTRDPTTPLARAAWETARHTPVNSWLGNIIMYAPTLW | 420 |
| PDB: 3Q0Z 23E | ELITSCSSNVSVAHDASGKRVYYLTRDPTTPLARAAWETARHTPVNSWLGNIIMYAPTLW | 420 |

**B**

##### Ligand Proximity to Thumb-1 Pocket: Amino Acids 421-480

| | | |
|---|---|---|
| PDB: 2BRK CMF | ARMILMTHFFSILLAQEQLEKALDCQIYGACYSIEPLDLPQIIERLHGLSAFSLHSYSPG | 480 |
| PDB: 2BRL POO | ARMILMTHFFSILLAQEQLEKALDCQIYGACYSIEPLDLPQIIERLHGLSAFSLHSYSPG | 480 |
| PDB: 2DXS JTP | ARMILMTHFFSILLAQEQLEKALDCQIYGACYSIEPLDLPQIIERLHGLSAFSLHSYSPG | 480 |
| PDB: 2XWY IB8 | ARMILMTHFFSILLAQEQLEKALDCQIYGACYSIEPLDLPQIIERLHGLSAFSLHSYSPG | 480 |
| PDB: 2WCX VGC | ARMILMTHFFSILLAQEQLEKALDCQIYGACYSIEPLDLPQIIERLHGLSAFSLHSYSPG | 480 |
| PDB: 3MWW BIW | ARMILMTHFFSILLAQEQLEKALDCQIYGACYSIEPLDLPQIIERLHGLSAFTLHSYSPG | 480 |
| PDB: 4DRU 0LN | ARMILMTHFFSILLAQEQLEKALDCQIYGACYSIEPLDLPQIIERLHGLSAFTLHSYSPG | 480 |
| PDB: 4GMC 1BI | ARMILMTHFFSILLAQEQLEKALDCQIYGACYSIEPLDLPQIIERLHGLSAFTLHSYSPG | 480 |
| PDB: 4NLD 2N7 | ARMILMTHFFSILLAQEQLEKALDCQIYGACYSIEPLDLPQIIQRLHGLSAFSLHSYSPG | 480 |
| PDB: 5TRK 23E | ARMILMTHFFSILLAQEQLEKALDCQIYGACYSIEPLDLPQIIQRLHGLSAFSLHSYSPG | 480 |
| PDB: 3Q0Z 23E | ARMILMTHFFSILLAQEQLEKALDCQIYGACYSIEPLDLPQIIQRLHGLSAFSLHSYSPG | 480 |

**C**

##### Ligand Proximity to Thumb-1 Pocket: Amino Acids 481-540

| | | |
|---|---|---|
| PDB: 2BRK CMF | EINRVASCLRKLGVPPLRVWRHRARSVRARLLSQGGRAATCGKYLFNWAVKTKLKLTPIP | 540 |
| PDB: 2BRL POO | EINRVASCLRKLGVPPLRVWRHRARSVRARLLSQGGRAATCGKYLFNWAVKTKLKLTPIP | 540 |
| PDB: 2DXS JTP | EINRVASCLRKLGVPPLRVWRHRARSVRARLLSQGGRAATCGKYLFNWAVKTKLKLTPIP | 540 |
| PDB: 2XWY IB8 | EINRVASCLRKLGVPPLRVWRHRARSVRARLLSQGGRAATCGKYLFNWAVKTKLKLTPIP | 540 |
| PDB: 2WCX VGC | EINRVASCLRKLGVPPLRVWRHRARSVRARLLSQGGRAATCGKYLFNWAVKTKLKLTPIP | 540 |
| PDB: 3MWW BIW | EINRVASCLRKLGVPPLRTWRHRARSVRAKLLSQGGRAATCGRYLFNWAVRTKLKLTPIP | 540 |
| PDB: 4DRU 0LN | EINRVASCLRKLGVPPLRTWRHRARSVRAKLLSQGGRAATCGRYLFNWAVRTKLKLTPIP | 540 |
| PDB: 4GMC 1BI | EINRVASCLRKLGVPPLRTWRHRARSVRAKLLSQGGRAATCGRYLFNWAVRTKLKLTPIP | 540 |
| PDB: 4NLD 2N7 | EINRVASCLRKLGVPPLRVWRHRARSVRARLLSQGGRAATCGKYLFNWAVRTKLKLTPIP | 540 |
| PDB: 5TRK 23E | EINRVASCLRKLGVPPLRVWRHRARSVRARLLSQGGRAATCGKYLFNWAVRTKLKLTPIP | 540 |
| PDB: 3Q0Z 23E | EINRVASCLRKLGVPPLRVWRHRARSVRARLLSQGGRAATCGKYLFNWAVRTKLKLTPIP | 540 |

***Supplementary FIG S1 (A-C)*** *Displays aligned amino acid residues of the HCV RdRp Thumb-1 pocket near respective ligands in PDB Structures, with PDB and ligand IDs listed. Residues comprising atomic coordinates closer than 5Å (yellow) and 3.5Å (red) to the ligand are labeled.*
